# Differential roles of frontoparietal regions in working memory and decision-making

**DOI:** 10.64898/2026.09.23.753872

**Authors:** Sophie E. Ack, Samantha M. Gray, Adam J.O. Dede, David King-Stephens, Kenneth D. Laxer, Ignacio Saez, Fady Girgis, Stephan U. Schuele, Joshua M. Rosenow, Eishi Asano, Robert T. Knight, Rodrigo M. Braga, Elizabeth L. Johnson

## Abstract

Working memory (WM) and decision-making (DM) are closely linked cognitive processes that rely on frontoparietal regions, yet their distinct and shared neural dynamics remain unclear. Using intracranial EEG recordings from 24 human subjects performing a delayed match-to-sample task, we examined high-frequency broadband (HFB) activity to dissociate WM- and DM-related processes with high spatiotemporal resolution. Principal component analysis of frontoparietal HFB activity revealed multiple temporally distinct patterns spanning stimulus presentation and delay phases. The most dominant component reflected sustained WM-related activity across task phases, consistent with persistent maintenance of internal representations. Additional components captured transient WM encoding across trial phases, as well as both stimulus-linked and ramping DM-related activity suggesting dual early and late stages of evaluative processing. These processes were distributed across frontoparietal regions but showed systematic differences in anatomical focus, with WM-related activity concentrated in dorsolateral prefrontal cortex (PFC) and DM-related activity concentrated in ventrolateral PFC, while inferior parietal cortex preferentially tracked the behavioral significance of incoming stimuli and superior parietal cortex tracked the appearance of new information that may require updating ongoing processing. Together, these findings demonstrate that WM and DM processes are separable yet interleaved over time within distributed cortical networks, revealing coordinated but functionally distinct contributions to goal-directed behavior.

## 1. Introduction

Working memory (WM) and decision-making (DM) are complex, higher-order cognitive functions that are intrinsically linked. WM enables the temporary storage and manipulation of information necessary for goal-directed behavior, while DM processes include evaluating incoming information against stored content to guide behavioral responses. Converging evidence from human neuroimaging, intracranial electroencephalography (iEEG), and non-human primate electrophysiology has established a distributed network largely composed of lateral prefrontal (PFC) and posterior parietal (PPC) cortices as central to WM processing (Baddeley, 2012; Buckner, 2003; Goldman-Rakic, 1995; Miller & Cohen, 2001). Activity within this network has been linked to multiple stages of WM, including stimulus encoding and maintenance of internal representations, and has also been implicated in DM evaluative processes, which may unfold concurrently with or subsequent to WM maintenance. Indeed, iEEG studies have demonstrated neural signals associated with stimulus evaluation and decision formation within frontoparietal cortices (Brosnan et al., 2020; Dürschmid et al., 2016; Liu & Pleskac, 2011; Rangarajan et al., 2020; Usher et al., 2013), which may be compatible with temporally overlapping WM processes. However, the relationship between WM- and DM-related processes and the distribution of their shared and distinct roles within frontoparietal regions remains undefined.

iEEG recordings present a means of investigating the neural dynamics underlying cognitive processes, providing sub-second temporal and centimeter spatial precision. High-frequency broadband (HFB; 70-150 Hz) activity recorded from human cortex closely tracks local neuronal population activity and is widely used to index task-related neural engagement (Lachaux et al., 2012; Lei et al., 2026; Manning et al., 2009). Previous iEEG studies have demonstrated robust HFB responses in frontal and parietal cortices during WM tasks (Johnson et al., 2020; Johnson et al., 2023), including sustained activity across delay periods that is consistent with the maintenance of internal representations (Haller et al., 2018; Rangarajan et al., 2020).

Traditional models of WM emphasize persistent neuronal activity as the mechanism of short-term storage during delay periods (Goldman-Rakic, 1995; Kamiński & Rutishauser, 2020; Miller & Cohen, 2001). However, recent work suggests that WM representations also exist in latent or “activity-silent” states that are not reflected in sustained neuronal activity (Baddeley, 2012; Barbosa et al., 2020; Hedayati et al., 2022; Miller et al., 2018; Sprague et al., 2016; Stokes, 2015; Wolff et al., 2015). In this framework, stored representations may instead be maintained through transient neural changes, rather than persistent activity, and reactivated when needed. Presentation of task-irrelevant stimuli has provided evidence supporting reactivation of otherwise latent stored representations during the delay period (Barbosa et al., 2020; Wolff et al., 2015; Wolff et al., 2017). Together, these findings suggest that WM processing involves dynamic transitions between active and latent states over time.

A major challenge in characterizing these dynamics is that neural activity recorded during WM tasks often reflects a mixture of distinct but overlapping processes, including possibly concurrent WM- and DM-related processes that can be difficult to parse. Many studies have focused on activity within individual cortical regions by averaging signals across electrodes that share anatomy but may be functionally distinct (Mercier et al., 2022), which can obscure parallel yet distinct signals. Identifying separable patterns of activity that correspond to WM and DM processes requires an analytical approach capable of capturing reliable, temporally structured patterns of activity across distributed neuronal populations. To accomplish this, we leveraged principal component analysis (PCA) to characterize responses across frontoparietal cortices by functional profile rather than anatomical segmentation. On half the trials, we further leveraged PCA time-locked to the presentation of a task-irrelevant stimulus to further characterize potentially active and latent processes associated with WM and DM.

We aimed to characterize how WM and DM processes are functionally organized across frontal and parietal cortices, and to establish the dynamic spatiotemporal evolution of these processes as reflected in HFB power throughout task performance. We used iEEG recordings from adult neurosurgical epilepsy patients with seizure- and artifact-free electrodes in lateral PFC and PPC during performance of a WM delayed match-to-sample (DMS) task. DMS paradigms are well-suited to characterize these processes because they require subjects to maintain a representation of previously observed stimuli and compare them with subsequently presented information during the test portion of the task to guide behavioral responses. We hypothesized that HFB activity would reveal dissociable spatial and temporal patterns associated with WM and DM, with distinct frontoparietal regions preferentially expressing each process and their relative contributions evolving over the course of the task. We applied PCA across segments of the WM-DMS task to examine task-related changes in HFB power and, specifically, how WM and DM processes evolve within distributed frontoparietal cortices. We also presented a brief task-irrelevant stimulus in order to disrupt ongoing processing and performed additional regional analyses to probe anatomical organization. Our findings, including temporally distinct early stimulus-related WM effects and ramping DM-related dynamics during delay, motivated subsequent analyses which revealed perturbation-evoked reactivation of latent WM representations and specialized organization of WM and DM processes across frontoparietal cortices.

## 2. Methods

### 2.1 Subjects

Intracranial EEG recordings were obtained from 24 subjects (M ± SD, 27.4 ± 10.8 years of age; 8 females) across 10 centers (California Pacific Medical Center; Ichan School of Medicine at Mount Sinai; Northwestern Memorial Hospital; Ohio State University; University of California, Davis; University of California, Irvine; University of California, San Diego; University of California, San Franciso; & Wayne State University), all undergoing presurgical monitoring for intractable epilepsy. Subjects were included based on above-chance task performance (inclusion criterion: <0.35 error in each condition, chance 0.5; (Johnson et al., 2023)) and non-pathological electrode coverage in lateral prefrontal and/or parietal grey matter. Demographic details are provided in Table 1. All subjects had normal/corrected-to-normal vision and hearing, no lesions evident on structural MRI or prior resections, and no other neurological or psychiatric diagnoses. All patients provided written informed consent in accordance with the Declaration of Helsinki as part of the research protocol approved by the Institutional Review Board at each institution.

**Table 1.** Subject demographics and electrode coverage. PFC, prefrontal cortex; PPC, posterior parietal cortex; sEEG, stereoelectroencephalograpy; ECoG, electrocorticography; R, right; L, left.

| <u>Subject</u> | <u>Type</u> | <u>Age</u> | <u>Sex</u> | <u>Seizure</u><br><u>Onset</u><br><u>Location</u> | <u>Trials (n)</u> |  |  | <u>Electrodes (n)</u> |  |  | <u>Responsive</u><br><u>Electrodes (n)</u> |  |  | <u>Reactivity (%)</u> |  |  |
| --- | --- | --- | --- | --- | --- | --- | --- | --- | --- | --- | --- | --- | --- | --- | --- | --- |
|  |  |  |  |  | No<br>Star | Star | Total | PFC | PPC | Total | PFC | PPC | Total | PFC | PPC | Total |
| 1 | sEEG | 15 | M | unknown | 63 | 59 | 122 | 13 | 0 | 13 | 8 | 0 | 8 | 61.5 |  | 61.5 |
| 2 | ECoG | 26 | M | seizures not<br>captured | 49 | 53 | 102 | 67 | 0 | 67 | 47 | 0 | 47 | 70.1 |  | 70.1 |
| 3 | ECoG | 31 | M | unknown | 58 | 59 | 117 | 10 | 22 | 32 | 3 | 3 | 6 | 30.0 | 13.6 | 18.8 |
| 4 | sEEG | 51 | F | unknown | 44 | 46 | 90 | 0 | 3 | 3 | 0 | 1 | 1 |  | 33.3 | 33.3 |
| 5 | sEEG | 27 | M | R frontal<br>temporal | 55 | 50 | 105 | 16 | 5 | 21 | 10 | 0 | 10 | 62.5 | 0.0 | 47.6 |
| 6 | sEEG | 28 | F | L temporal | 57 | 59 | 116 | 0 | 10 | 10 | 0 | 0 | 0 |  | 0.0 | 0.0 |
| 7 | sEEG | 38 | F | unknown | 61 | 53 | 114 | 25 | 0 | 25 | 5 | 0 | 5 | 20.0 |  | 20.0 |
| 8 | sEEG | 30 | M | unknown | 61 | 61 | 122 | 0 | 5 | 5 | 0 | 3 | 3 |  | 60.0 | 60.0 |
| 9 | ECoG | 16 | F | seizures not<br>captured | 52 | 52 | 104 | 34 | 11 | 45 | 16 | 7 | 23 | 47.1 | 63.6 | 51.1 |
| 10 | sEEG | 27 | M | bilateral<br>temporal | 54 | 50 | 104 | 13 | 0 | 13 | 12 | 0 | 12 | 92.3 |  | 92.3 |
| 11 | sEEG | 26 | M | R temporal | 54 | 48 | 102 | 20 | 0 | 20 | 2 | 0 | 2 | 10.0 |  | 10.0 |
| 12 | sEEG | 25 | F | R<br>supplemental<br>motor area | 56 | 59 | 115 | 24 | 0 | 24 | 11 | 0 | 11 | 45.8 |  | 45.8 |
| 13 | sEEG | 21 | F | bilateral<br>temporal | 52 | 49 | 101 | 9 | 7 | 16 | 6 | 0 | 6 | 66.7 | 0.0 | 37.5 |
| 14 | sEEG | 41 | M | bilateral<br>temporal | 57 | 57 | 114 | 16 | 0 | 16 | 2 | 0 | 2 | 12.5 |  | 12.5 |
| 15 | sEEG | 32 | M | unknown | 57 | 61 | 118 | 5 | 10 | 15 | 1 | 2 | 3 | 20.0 | 20.0 | 20.0 |
| 16 | sEEG | 54 | M | unknown | 59 | 54 | 113 | 6 | 0 | 6 | 1 | 0 | 1 | 16.7 |  | 16.7 |
| 17 | sEEG | 23 | M | unknown | 52 | 54 | 106 | 14 | 7 | 21 | 3 | 0 | 3 | 21.4 | 0.0 | 14.3 |
| 18 | sEEG | 32 | M | R temporal | 49 | 55 | 104 | 33 | 0 | 33 | 8 | 0 | 8 | 24.2 |  | 24.2 |
| 19 | sEEG | 32 | F | bilateral<br>frontal<br>temporal | 37 | 30 | 67 | 22 | 0 | 22 | 9 | 0 | 9 | 40.9 |  | 40.9 |
| 20 | ECoG +<br>sEEG | 23 | M | L temporal<br>parietal | 52 | 53 | 105 | 0 | 11 | 11 | 0 | 0 | 0 |  | 0.0 | 0.0 |
| 21 | sEEG | 13 | F | L insula | 56 | 51 | 107 | 0 | 17 | 17 | 0 | 10 | 10 |  | 58.8 | 58.8 |
| 22 | sEEG | 14 | M | L temporal | 11 | 19 | 30 | 15 | 42 | 57 | 9 | 20 | 29 | 60.0 | 47.6 | 50.9 |
| 23 | sEEG | 15 | M | L insula | 53 | 53 | 106 | 20 | 0 | 20 | 9 | 3 | 12 | 45.0 |  | 60.0 |
| 24 | sEEG | 17 | M | unknown | 49 | 53 | 102 | 0 | 11 | 11 | 0 | 7 | 7 |  | 63.6 | 63.6 |
| **Total** |  |  |  |  |  |  |  | **362** | **161** | **523** | **162** | **56** | **218** | **44.8** | **34.8** | **41.7** |

### 2.2 Delayed match-to-sample task

Subjects performed a DMS task in which they compared three colored shapes in specific spatiotemporal positions in a test sequence to their internal representation of a sample sequence (Figure 1) (Cross et al., 2025; Gray et al., 2026; Shi et al., 2025; Yarbrough et al., 2025). On each trial, subjects focused on a central fixation crosshair and observed the 1.5-s sample sequence of shapes (0.25-s stimulus, 0.25-s interstimulus interval [ISI]) followed by a 1.75-s delay, then a test sequence with the same timing of stimulus presentation and delay. In half of all trials, we briefly flashed a star for 0.1-s at a randomly jittered time ranging from 0.75-0.9-s from the onset of the post-sample and post-test delays, which subjects were told to ignore. The star was a task-irrelevant stimulus designed to disrupt ongoing processing and elicit measurable responses related to otherwise latent WM representations (Wolff et al., 2015). Specifically, Wolff et al. (2015) found that presenting such a stimulus during the delay elicited a neural response which carried significant information about the contents of information held in working memory. After each post-test delay, subjects reported by self-paced mouse click whether the sample and test sequences were matched or mismatched. Mismatched sequences always included mismatches in two of the three shapes, in one of three dimensions: shape identity, spatial orientation, or temporal order. In effect, this design positioned only shapes 1 and 2 as strictly necessary for task performance, although the entire sequence is considered in the sole decision for each trial. Following eight practice trials, subjects completed 128 trials with a break every 16 trials. Test conditions and star trials were presented randomly and evenly counterbalanced across sets of 16 trials. Only correct trials were included in subsequent analyses. The task was programmed in MATLAB (MathWorks Inc., Natick, MA) with the Psychtoolbox-3 software extension.

**Figure 1.**
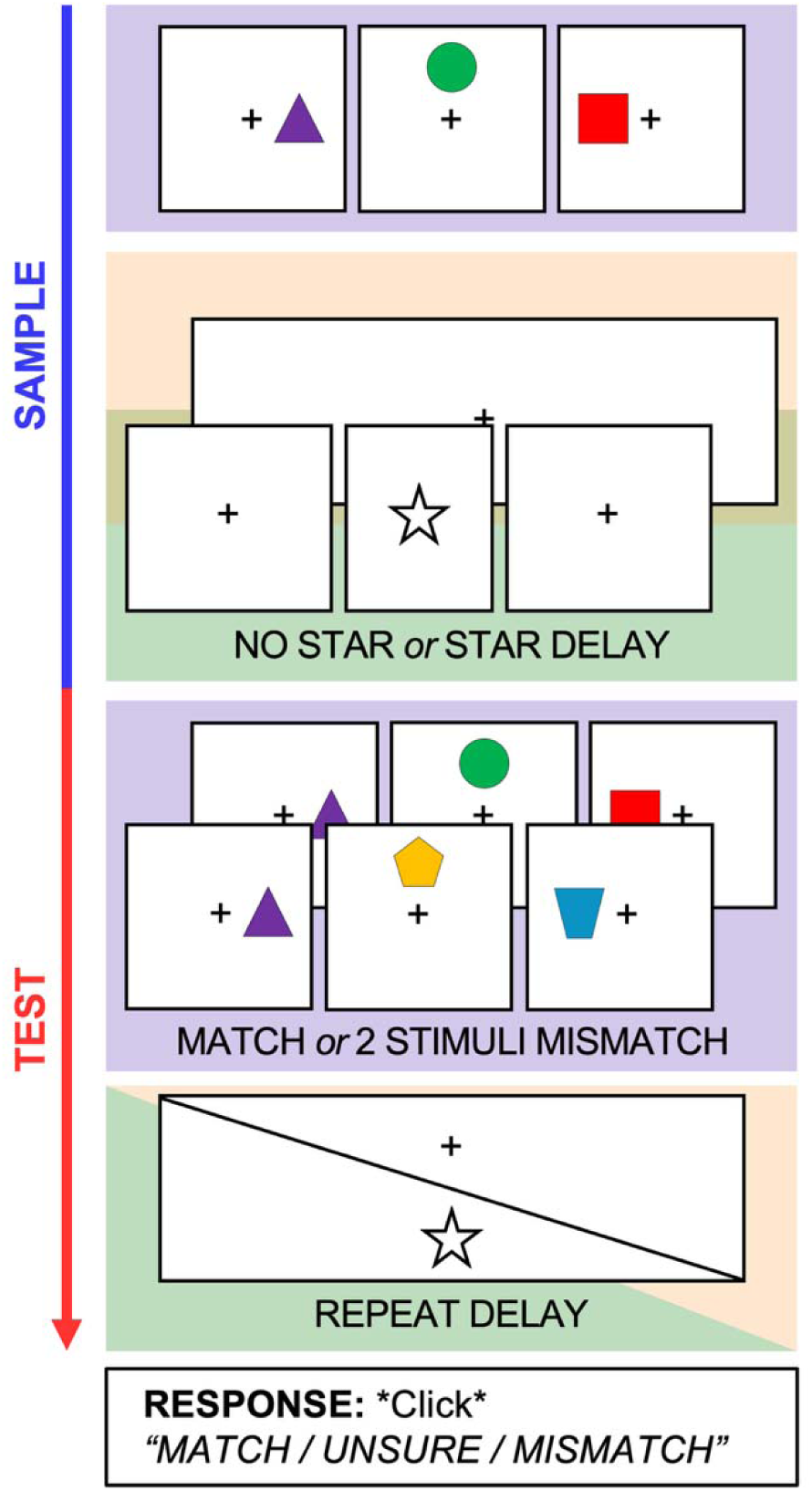
Working memory delayed match-to-sample task paradigm. On each trial, subjects viewed a sample sequence of three shapes in specific spatiotemporal positions over a 1.5-s period (0.25 s per stimulus, separated by interstimulus interval [ISI] periods of 0.25 s each; purple, top), followed by a 1.75-s post-sample delay. An errant star stimulus was briefly flashed at a randomly jittered time during the delay (0.1 s duration, onsetting 0.75 to 0.9 s into the delay) in half of all trials (star delay, orange), which participants were told to ignore, versus no interruption to fixation during the other half of all trials (no-star delay, green). After the delay, a test sequence was presented with identical timing to the sample sequence (purple, middle), and either matched the sample sequence exactly (above) or contained 2 of 3 mismatched stimuli in either identity, spatial, or temporal dimensions (below, with identity mismatch illustrated). The same type of post-sample delay (i.e. no star or star) occurred post-test, with identical star timing if presented. At the end of each trial, subjects reported if the sample and test sequences were matched or mismatched by self-paced mouse click. Resulting trial segments for sample vs. test analysis were stimulus presentation (purple), no-star delay (orange), and star delay (green).

### 2.3 Electrode placement and localization to frontal or parietal cortices

Macro-electrodes were surgically implanted for extra-operative recording based solely on the clinical needs of each patient. These electrodes were placed subdurally in 64-channel grids and 4- and 8-channel strips with 10-mm spacing (ECoG n = 4), and/or stereotactically in 8-, 10-, and 12-channel tracks with 5-mm spacing (sEEG n = 21). Anatomical locations were determined by co-registering post-implantation computed tomography (CT) coordinates to pre-operative magnetic resonance (MR) images, as implemented in FieldTrip (Stolk et al., 2018). Electrodes were localized in native space based on visual inspection of individual anatomy to manually select frontal and parietal grey matter contacts. We localized electrodes to subregions of the lateral prefrontal cortex (inferior frontal gyrus, IFG; middle frontal gyrus, MFG; and superior frontal gyrus, SFG) and posterior parietal cortex (angular gyrus, AG; supramarginal gyrus, SMG; and superior parietal lobule, SPL) using major sulcal landmarks. They were transformed into standard Montreal Neurological Institute (MNI) space for visualization across subjects (Figure 2).

**Figure 2.**
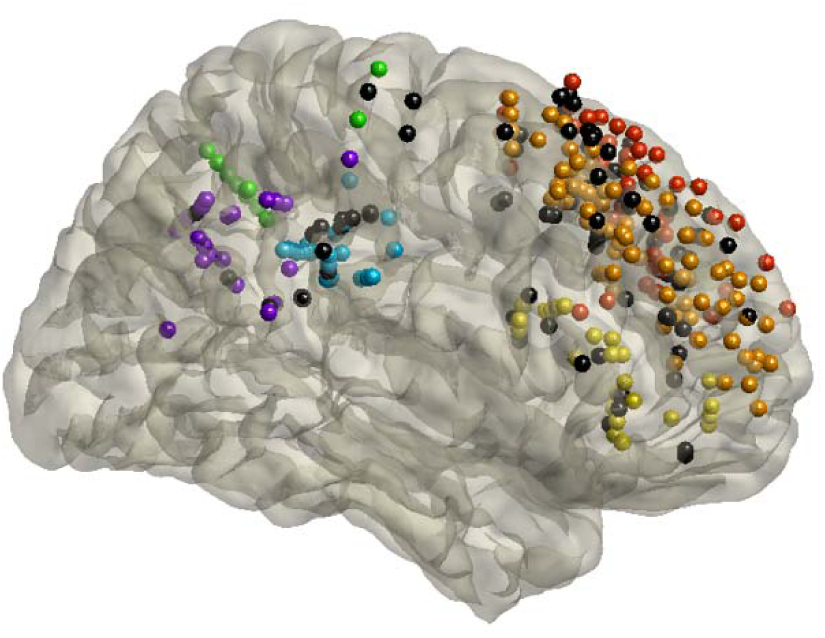
Electrode locations. Implanted electrodes across all subjects in the lateral frontal and parietal cortices shown on a Montreal Neurological Institute (MNI) template brain, in lateral perspective and mirrored across hemispheres. Responsive electrodes included in the analyses in the frontal cortex are shown in warm colors: superior frontal gyrus (SFG) in red, middle frontal gyrus (MFG) in orange, and interior frontal gyrus (IFG) in yellow. Responsive electrodes included in the analyses in the parietal cortex are shown in cool colors: superior parietal lobule (SPL) in green, supramarginal gyrus (SMG) in blue, and angular gyrus (AG) in purple. Subregion membership in frontal and parietal cortices was defined by major sulcal boundaries within individuals. Electrodes shown in black were nonresponsive electrodes that were excluded from analyses (as described in Methods).

### 2.4 Intracranial EEG preprocessing

Electrophysiological data were acquired using 128/256-channel Nihon Kohden (Nihon Kohden Corp., Tokyo, Japan) or Natus (Natus Medical Inc., Middleton, WI, USA) recording systems, sampled at 5 kHz, 1 kHz, or 512 Hz. Data sampled at 5 kHz were resampled to 1 kHz offline. Raw data were filtered with 0.1-Hz high-pass and, if sampled at 1 kHz, 300-Hz low-pass finite impulse response filters, and 60-Hz line noise harmonics were removed using discrete Fourier transform. Continuous data were demeaned, epoched into 8.5-s trials (−1 s from the onset of the sample sequence to +1s from the offset of the post-test delay), and manually inspected blind to electrode locations and experimental task parameters. Electrodes in seizure onset zones and electrodes and epochs displaying epileptiform activity or artifactual signal (from poor contact, machine noise, etc.) were excluded, ensuring that data considered for analysis represent healthy tissue (Rossini et al., 2017). Neighboring electrodes within the same anatomical structure were then bipolar montage re-referenced using consistent conventions (ECoG strips, anterior – posterior; sEEG, deep – surface) (Cross et al., 2025; Gray et al., 2026; Johnson et al., 2018; Johnson et al., 2019; Johnson et al., 2023; Kam et al., 2021; Shi et al., 2025). For ECoG grids, electrodes were re-referenced to neighboring electrodes on a row-by-row basis. Re-referenced data were manually re-inspected to reject any trials with residual noise, resulting in 103.6 ± 19.4 (52.0 ± 10.4 no-star; 51.6 ± 9.4 star; M ± SD) seizure- and artifact-free, correct trials analyzed per subject. All data preprocessing and analysis were performed using custom MATLAB functions, with the EEGLAB (Delorme & Makeig, 2004) and Fieldtrip (Oostenveld et al., 2011) toolboxes. Electrodes within regions of interest were considered for subsequent analyses.

### 2.5 High-frequency broadband analyses

#### 2.5.1 Computation of HFB activity

Power was computed from 70 to 150 Hz using a multitapering approach (Mitra & Pesaran, 1999) in steps of 10 Hz (i.e., 70–80, 80–90… 140–150 Hz). Data segments were zero-padded to the next power of 2 to minimize filter-induced artifacts, and the multitaper frequency spectrum was calculated by sliding a 300-ms window in 5-ms increments. An output resolution of 5 ms was used to equate the time axis across datasets sampled at 1 kHz and 512 Hz while preserving temporal precision. HFB activity was analyzed using statistical bootstrapping (Flinker et al., 2015; Haller et al., 2018; Johnson et al., 2018; Johnson et al., 2023; Kam et al., 2021; Kam et al., 2019; Rangarajan et al., 2020). For each electrode and frequency, baseline (−250 to −50 ms from the onset of the sample sequence) raw power values were pooled into a single time series, from which 200 datapoints were randomly selected and averaged (Johnson et al., 2023; Kam et al., 2021). This step was repeated 1000 times to create bootstrapped distributions of baseline data. Post-stimulus (0 to 6.5 s from the onset of the sample sequence) raw power data were z-scored on the baseline distributions and then averaged across frequencies. This procedure adjusts the power outputs to correct for 1/f power scaling and reveals significant HFB activity that is elicited by the WM task.

Electrodes were considered responsive if mean HFB activity was above threshold (z = 1.96, equivalent to α = 0.05, two tailed) for at least 10 consecutive timepoints (i.e., 50 ms) across trials. This duration criterion exceeds the minimum number of consecutive timepoints requisite for a time series to be considered significantly different from zero (Flinker et al., 2015; Guthrie & Buchwald, 1991; Haller et al., 2018; Johnson et al., 2023). This analysis is similar to methods used in single-unit neurophysiology and is naïve regarding electrode location and task condition. Across hemispheres, 44.8% (162 of 362) of frontal electrodes and 34.8% (56 of 161) of parietal electrodes met task-responsive criteria. These proportions are consistent with previous studies in the field (Haller et al., 2018; Johnson et al., 2023). The locations of these electrodes are shown in Figure 2, and per-subject counts are provided in Table 1.

#### 2.5.2 Epoch definition

Following HFB computation, data from each trial were further epoched into the three segments of the task (Figure 1). For all trials, stimulus presentation was defined from onset of the first shape in each sequence (0-s in sample, 3.25-s in test) to offset of the third ISI (1.5-s in sample, 4.75-s in test; 1.5-s duration). In no-star trials, the delay period was defined from the end of stimulus presentation (1.5-s in sample, 4.75-s in test) to the end of the delay (3.25-s in sample, 6.5-s in test; 1.75-s duration). In star trials, the delay period data were realigned with the zero point at the onset of the star and cut into 1.6-s (-0.75 to 0.85-s from star onset) epochs, to account for the jitter in star presentation. All subsequent analyses were separately applied to the stimulus presentation, no-star delay, and star delay segments.

### 2.6 Modulation of HFB by cognitive demand

#### 2.6.1 Cohen’s d for effect size

To identify neural activity preferentially associated with WM and DM, we examined modulation of HFB activity by cognitive demand by comparing activity at matching timepoints in sample and test. Cohen’s d was computed to measure the effect size for each channel at each timepoint, using the following equation:

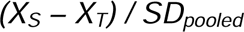

where *X_S_* is the sample mean across trials, *X_T_*is the test mean across trials, and *SD_pooled_* is the pooled standard deviation across both sample and test (Cohen, 1988). Positive effect sizes indicate sample-specific WM processes associated with encoding and maintenance, while negative effect sizes indicate test-specific DM processes associated with retrieval and evaluative processing.

#### 2.6.2 Principal component analysis

We performed principal component analysis (PCA) on the derived effect sizes for the stimulus presentation, no-star delay, and star delay epochs to identify separable and temporally consistent patterns of HFB activity accounting for the most variance within responsive electrodes across subjects. All responsive electrodes from both frontal and parietal regions were included as a single population in the PCA. We used a clear flattening in the variance explained by the subsequent principal component (PC) to select PCs for further analysis, quantitatively including PCs until there was a less than 1% difference in variance explained by the next PC (Bolt et al., 2022; Chen et al., 2023; Dürschmid et al., 2016; Dürschmid et al., 2019). This resulted in four PCs per epoch. To define patterns of activity from the most representative electrodes, we extracted electrodes that were highly loaded into each of these PCs, defined as electrodes that reached the 75^th^ percentile of loadings (henceforth referred to as PC-loading electrodes: PC1-loading electrodes, PC2-loading electrodes, etc.). Subsequent statistical modeling was performed on HFB timeseries data (rather than effect sizes) in these PCA-defined sets of electrodes. These procedures are consistent with those employed in several iEEG studies using different task paradigms (Dürschmid et al., 2016; Dürschmid et al., 2019; Kam et al., 2021).

#### 2.6.3 Subregion anatomical analysis

In addition to grouping electrodes by function through PCA, we repeated analyses of HFB timeseries data in electrodes defined by anatomical subregion to test the apparent spatial effects of WM and DM observed in our PCA findings, which suggested localized processes or specialized subregions. Responsive electrodes in each of the previously defined frontal and parietal subregions (IFG, MFG, SFG; AG, SMG, SPL) were selected as separate populations.

### 2.7 Group-level statistics

#### 2.7.1 Linear mixed-effects models

All reported results are based on linear mixed-effects (LME) models of single-electrode differences between sample and test, with subjects and nested electrodes as random intercepts (Yu et al., 2022). Subjects and nested electrodes were modeled as random to indicate repeated measures and account for sources of inter-individual variability including electrode counts and anatomical sampling (Johnson & Knight, 2023). Because signal components varied across time, LME models were applied at every timepoint independently.

#### 2.7.2 Cluster correction

We used a cluster-based permutation test to correct for multiple comparisons and depict the time course of significant differences between sample and test (Maris & Oostenveld, 2007). At each timepoint, the per-channel HFB power at sample and test was compared to a null distribution, estimated by randomly shuffling sample and test conditions for each trial and refitting the LME model, repeated 1000 times (Dede et al., 2025). The *p*-values of the observed clusters were calculated as the proportion of random partitions that yielded a larger effect than the observed experimental effect, thresholded at an alpha of 0.05 (Kam et al., 2021; Spaak et al., 2016). Resulting one-sided *p-*values were doubled for a two-tailed test (Spaak et al., 2016). Therefore, the smallest *p*-value possible in our analysis is 0.002.

## 3. Results

### 3.1 Behavioral performance

We evaluated task performance by calculating behavioral accuracy rate per subject as the hit rate (i.e., proportion of match trials that were correctly identified as match) minus the false alarm rate (proportion of mismatched trials that were incorrectly identified as match). This procedure defines chance accuracy as zero and corrects for differences in trial counts between conditions as well as an individual’s tendency to respond match/mismatch (Snodgrass & Corwin, 1988). We confirmed that iEEG patients performed well on the task (accuracy rate M ± SD: 0.82 ± 0.14; one-sample t(24) = 27.82, p < 0.001). Furthermore, accuracy rate did not differ from that of healthy controls (accuracy rate M ± SD: 0.85 ± 0.12; one-sample t(35) = 40.35, p < 0.001) (Shi et al., 2025; Yarbrough et al., 2025) as determined by Bayes Factor analysis (BF_10_ = 0.357) (Rouder & Morey, 2012). Additionally, neither accuracy rate (paired t-stat = 0.378, p = 0.709; BF_10_ = 0.292) nor reaction time (paired t-stat = 1.218, p = 0.236; BF_10_ = 0.432) differed between star (accuracy rate M ± SD: 0.85 ± 0.11; RT M ± SD: 2.14 ± 0.49 sec) and no star trials (accuracy rate M ± SD: 0.86 ± 0.09; RT M ± SD: 2.48 ± 1.59 sec). The absence of behavioral effects indicates that presentation of the task-irrelevant star did not measurably alter task performance, allowing neural differences between conditions to be interpreted independently of changes in behavioral demand, and supporting use of the star as a perturbation for probing latent WM and DM processes.

### 3.2 Working memory and decision-making effects

We identified dissociable patterns of activity associated with WM and DM by comparing neural responses during the sample and test phases of each trial segment of the DMS task, which were matched in sensory and temporal features but differed in their relative demands on memory maintenance and decision formation. To probe patterns of activity associated with WM, we identified PCs with significant clusters of positive effects (sample > test). To probe patterns of activity associated with DM, we identified PCs with significant clusters of negative effects (sample < test). Positive effects indicate sample-specific WM processes associated with encoding and maintenance, while negative effects indicate test-specific DM processes associated with recall and evaluative processing.

#### 3.2.1 Working memory encoding and maintenance effects are sustained for trial duration

The first PC was strongly implicated in WM in each of the three trial segments (i.e., stimulus presentation, no-star delay, star delay). Sample HFB power was elevated relative to test throughout all trial segments, which was sustained for the complete duration of stimulus presentation (0.000-1.500 s from shape 1 onset, *p* = 0.002, Figure 3A), no-star delay (1.500-3.250 s from delay onset, *p* = 0.002, Figure 3B), and star delay (-0.750-0.850 s from star onset, *p =* 0.002, Figure 3C). Thus, across all task epochs, the dominant component consistently reflected persistent WM-related activity.

**Figure 3.**
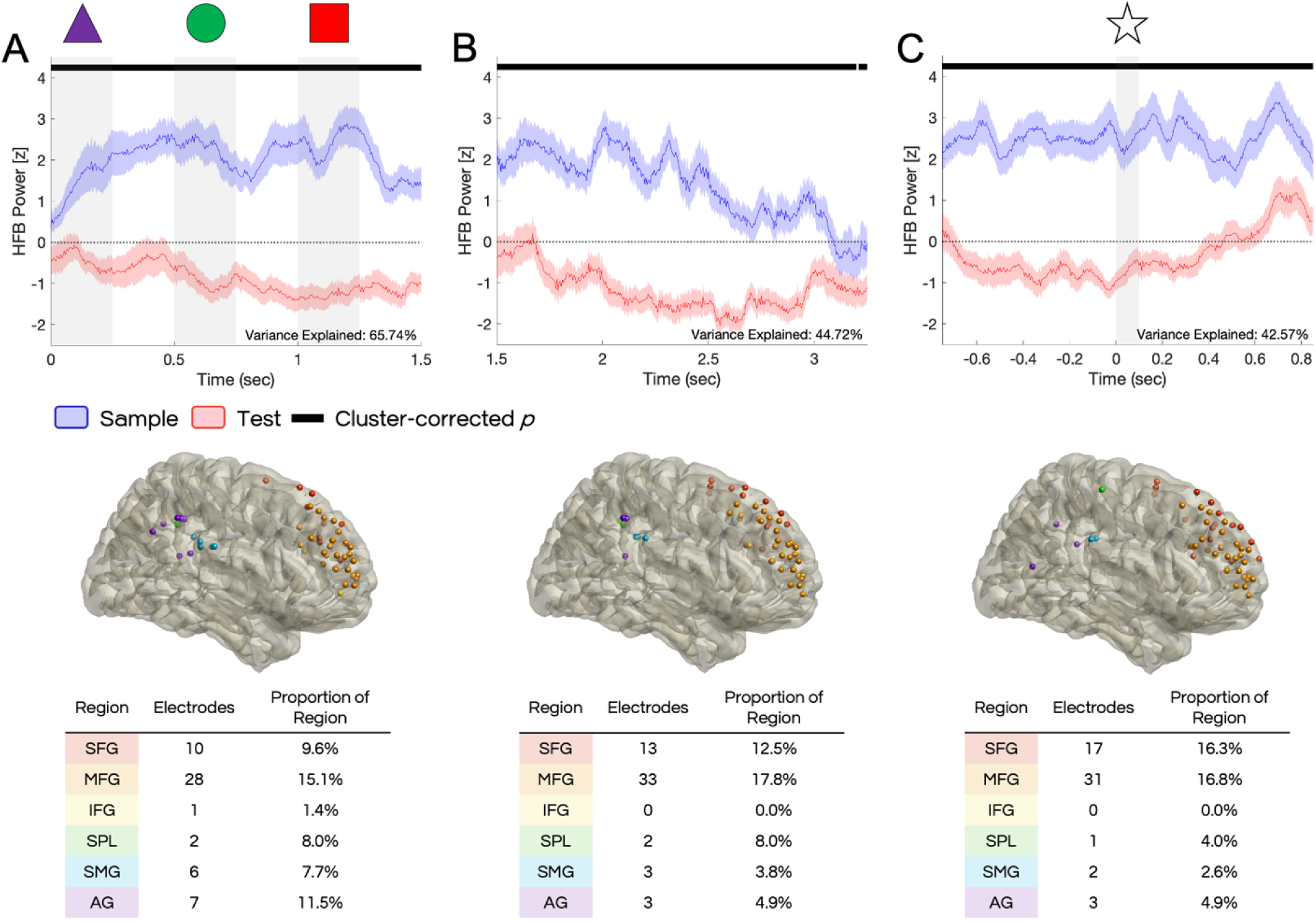
Working memory encoding and maintenance effects captured in PC1. Mean HFB timeseries from and locations of electrodes highly loading (75^th^ percentile of loadings) into the first principal component of each trial segment: (A) stimulus presentation, (B) no-star delay, and (C) star delay. Top: Mean z-scored HFB traces, with standard error shown in shaded area, for sample (blue) vs. test (red). Activity was consistently higher during sample than test in all trial segments. Black horizontal line indicates time points of significance by cluster-corrected *p*-value. Vertical grey shading represents duration of stimuli, with relevant stimulus represented above. Middle: PC-loading electrodes shown on an MNI template brain, in lateral perspective and mirrored across hemispheres. Bottom: breakdown of electrodes by subregion, by number and relative sampling compared to cohort electrode coverage in each region.

The first PCs represent the largest share of variance explained between sample and test HFB activity: stimulus presentation, 65.74%; no-star delay, 44.72%; star delay, 42.57%. The pattern of elevated sample activity was constant and unaffected by the presence of visual stimuli, albeit with a gradual decline in activity during the uninterrupted (no-star) delay. PC1-loading electrodes were located in both frontal and parietal regions in all three trial segments, with the majority in PFC. Proportional to cohort electrode coverage, MFG was the most represented (stimulus presentation: 28 electrodes, 15.1% of electrodes in region; no-star delay: 31 electrodes, 17.8% of electrodes in region; star delay: 33 electrodes, 16.8% of electrodes in region; Figure 3, bottom). In contrast, there was negligible representation of PC1-loading electrodes in IFG (stimulus presentation: 1 electrode, 1.4% of electrodes in region; no-star delay: 0 electrodes, 0.0% of electrodes in region; star delay: 0 electrodes, 0.0% of electrodes in region). These results support persistent, active WM maintenance throughout the task that appears selective to dorsal frontal regions including SFG and MFG.

#### 3.2.2 Neural activity tracks salient stimuli during sample stimulus presentation

Because mismatch trials always contain two mismatched shapes, shapes 1 and 2 of the sequence are most behaviorally relevant to completing the task since a mismatched shape appears by shape 2 in every mismatched trial. Subjects could ignore shape 3 during sample and test without impairing their behavioral results. This distinction between salient (shapes 1 and 2) and task irrelevant stimulus 3 was apparent in our PCA results.

PC2 of stimulus presentation demonstrated that WM-related activity preferentially tracked the behaviorally relevant portion of the stimulus sequence. Sample activity was elevated during the “salient period” of the first two shapes, from the onset of shape 1 until the offset of shape 2, after which sample activity returned to baseline (Figure 4A). The sample greater than test effect was significant for 475 ms, beginning 30 ms into shape 1 and ending shortly after the onset of shape 2 (0.030-0.505 s from shape 1 onset, *p =* 0.002). While the difference between sample and test during shape 2 was nonsignificant, this reflected increasing test activity rather than decreasing sample activity, which remained elevated above baseline through shape 2 before returning to baseline after its offset (see DM section to follow). This PC explained 8.19% of variance and had notably greater representation of IFG (4 electrodes, 5.6%) and inferior parietal regions (SMG: 7 electrodes, 9.0%; AG: 8 electrodes, 13.1%) than the WM effect-dominated PC1. PFC electrodes were also collectively more posterior than those in PC1.

**Figure 4.**
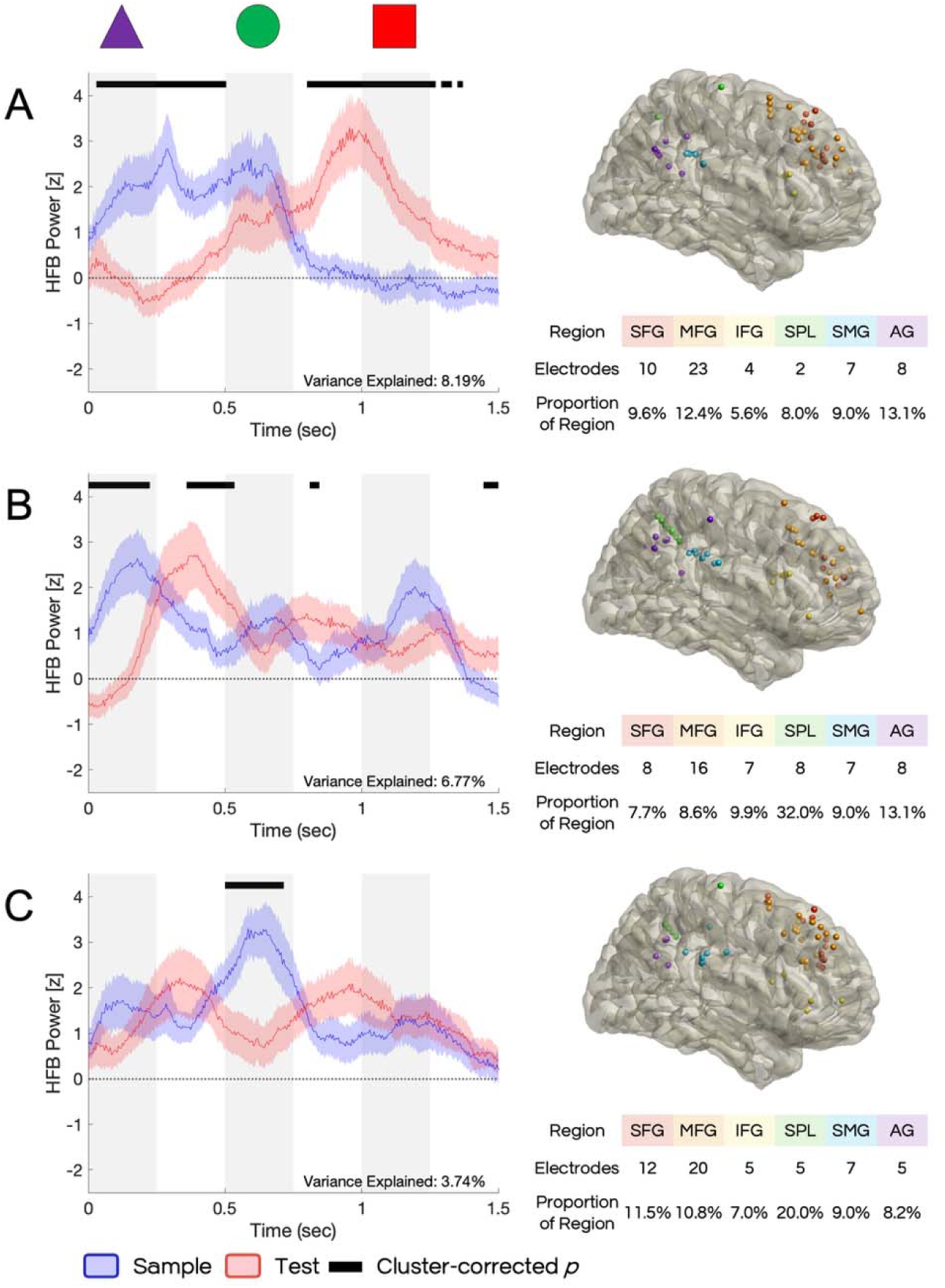
Effects associated with salient stimuli and early deliberative processing during stimulus presentation PC2-4. Mean HFB timeseries from and locations of electrodes highly loading (75^th^ percentile of loadings) into the second (A), third (B), and fourth (C) PCs of stimulus presentation. Left: Mean z-scored HFB traces, with standard error shown in shaded area, for sample (blue) vs. test (red). Black horizontal line indicates time points of significance by cluster-corrected *p*-value. Vertical grey shading represents duration of stimuli, with relevant stimulus represented above. Right, above: PC-loading electrodes shown on an MNI template brain, in lateral perspective and mirrored across hemispheres. Right, below: breakdown of electrodes by subregion, by number and relative sampling compared to cohort electrode coverage in each region.

Two additional components corroborated this salience-related WM effect by isolating transient responses to the presentation of shapes 1 and 2, respectively. PC3 captured elevated sample activity during shape 1. The significant cluster occurred over the first 225 ms of the trial, ending shortly before offset of the first shape at 250 ms (0.000-0.225 s from shape 1 onset, *p =* 0.002, Figure 4B). PC4 similarly isolated a significant sample effect during shape 2, lasting for 215 ms beginning with the onset of the second shape and ending shortly before its offset (0.500-0.715 s, *p* = 0.002, Figure 4C). These PCs explained 6.77% and 3.74% of variance, and a greater proportion of PC3- and PC4-loading electrodes were located in PPC, particularly in SPL (PC3: 32.0% SPL, 13.1% AG, 9.0% SMG; PC4: 20% SPL, 8.2% AG, 9.0% SMG) compared to stimulus presentation PC1. Similar to PC2, PFC electrodes were located more posteriorly.

Together, these results indicate that WM-related activity preferentially tracked behaviorally relevant stimuli. PC2 captured this effect across the salient portion of the sequence, while PC3 and PC4 further captured transient activity associated with shapes 1 and 2 individually. No significant WM effects were observed during the irrelevant third shape in any component.

#### 3.2.3 Stimulus DM effects are separable from WM salience detection

DM effects during stimulus presentation reflected both progressive evaluation of the stimulus sequence and transient processing associated with individual stimuli. PC2 revealed climbing increases in test activity with each stimulus, suggesting an accumulative DM process (Figure 4A). The activity began to increase during the first ISI and peaked around the onset of shape 3, with significance sustained for most of the second ISI and the entirety of shape 3 (0.800-1.370 s from shape 1 onset, *p* = 0.002). The activity peaked after shape 2 and began decreasing upon presentation of shape 3, further corroborating the distinction between salient and irrelevant stimuli. This trajectory may indicate an early match/mismatch deliberation accumulating through the behaviorally relevant portion of the sequence.

Individual test greater than sample clusters were observed during the ISI after each of the three shapes in PC3, with the highest power associated with shape 1 and the lowest power with shape 3 (Figure 4B). The first ISI cluster was also the longest, lasting for 175 ms and ending shortly after the onset of shape 2 (0.360-0.535 s from shape 1 onset, *p* = 0.002). The second and third ISI clusters were shorter, 35 ms and 55 ms respectively (0.810-0.845 s, *p* = 0.002; 1.445-1.500 s, *p* = 0.002). The delayed onsets of these stimulus-associated DM effects in PC3 (0.110 s from shape 1 offset, 0.060 s from shape 2 offset, 0.195 s from shape 3 offset) compared to the matched WM-effect for each shape may indicate additional processing time needed to integrate incoming test stimuli into a sequence for comparison with internal representations of the sample. Test effects did not reach significance in PC4 (Figure 4C).

Compared to PC1, which only captured WM effects, PC2- to PC-4-loading electrodes were much more prevalent in IFG (PC2: 4 electrodes, 5.6%; PC3: 7 electrodes, 9.9%; PC4: 5 electrodes, 7.0%), further suggesting IFG may play a uniquely DM-related role.

Together, these findings demonstrate that stimulus-related DM processing occurred at multiple temporal scales: PC2 captured a progressive evaluative process unfolding across the stimulus sequence, whereas PC3 isolated transient post-stimulus evaluation following individual stimuli. Both converged so that decision-related activity emerged before completion of the test sequence and preferentially tracked its behaviorally relevance.

#### 3.2.4 Delay period contains sustained DM effects

Additional DM effects were observed during the no-star delay period, indicating a secondary and prolonged evaluative process beyond the duration of the stimulus sequence. PC2 reveals inverse trajectories of activity during sample and test (Figure 5A). Where sample activity was elevated immediately following the end of stimulus presentation and gradually decreased below baseline through the delay, test activity began below baseline and climbed through the delay period. These trajectories intersected at baseline about 2 seconds into the overall trial, with significant WM effects before (sample > test 1.500-1.820 s from shape 1 onset, *p* = 0.002) and significant DM effects after (test > sample 2.305-3.250 s, *p* = 0.002). Viewed in the context of the full trial, these opposing trajectories suggest a natural temporal progression from diminishing WM-related activity during the sample delay to a gradually increasing deliberative DM process during the test delay that unfolds over a longer timescale than the early DM effects observed during stimulus presentation (see Figure 4).

**Figure 5.**
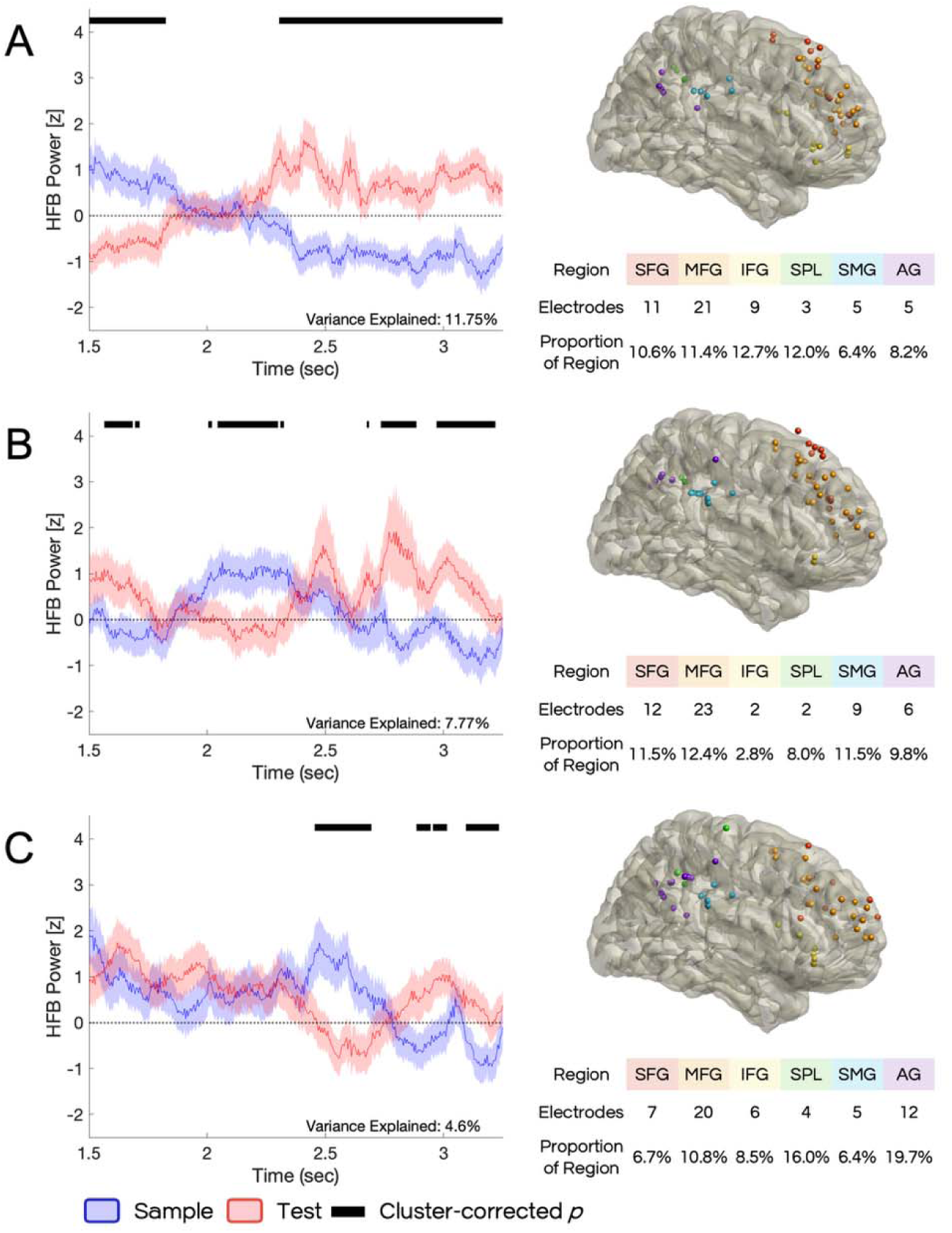
Delay decision-making effects in PC2-4. Mean HFB timeseries from and locations of electrodes highly loading (75^th^ percentile of loadings) into the second (A), third (B), and fourth (C) PCs of the no-star delay. Left: Mean z-scored HFB traces, with standard error shown in shaded area, for sample (blue) vs. test (red). Black horizontal line indicates time points of significance by cluster-corrected *p*-value. Vertical grey shading represents duration of stimuli, with relevant stimulus represented above. Right, above: PC-loading electrodes shown on an MNI template brain, in lateral perspective and mirrored across hemispheres. Right, below: breakdown of electrodes by subregion, by number and relative sampling compared to cohort electrode coverage in each region.

PC3 and PC4 indicated additional transient WM and DM effects during the no-star delay. In PC3, there were significant DM effects at the start and end of the delay (1.565-1.715 s from shape 1 onset, p = 0.002; 2.675-3.22 s, *p* = 0.006), with significant WM effects in the interim (2.005-2.325 s, *p* = 0.002; Figure 5B). In PC4, there was a WM effect (2.455-2.695 s from shape 1 onset, *p* = 0.002) followed by DM effects (2.885-3.235 s, *p* = 0.002), again intersecting at baseline (Figure 5C).

The anatomical distribution of these components paralleled that observed during stimulus presentation. Consistent with electrode distributions during stimulus presentation, PC2- to PC4-loading electrodes were prevalent in IFG (PC2: 9 electrodes, 12.7%; PC3: 2 electrodes, 2.8%; PC4: 6 electrodes, 8.5%) where PC1-loading electrodes contained none. Additionally, they contained a greater proportion of electrodes in PPC, particularly in SMG (PC2: 5 electrodes, 6.4%; PC3: 9 electrodes, 11.5%; PC4: 5 electrodes, 6.4%) and AG (PC2: 5 electrodes, 8.2%; PC3: 6 electrodes, 9.8%; PC4: 12 electrodes, 19.7%), than PC1. These PCs explained 11.75%, 7.77%, and 4.6% of the variance in the no-star delay period, respectively. All no-star delay PCs point to a handoff between WM and DM processing, with increases in DM temporally associated with decreases in WM. We next probed this antagonistic dynamic by analyzing the delay with the interruptive star stimulus, capitalizing on its ability to perturb ongoing processing and elicit measurable responses related to otherwise latent WM representations (Wolff et al., 2015).

### 3.3 Star interruption

Having observed the patterns of activity present during natural task performance without interference, we leveraged analysis of the star-interrupted delay period to probe potential latent or “activity silent” processes (Wolff et al., 2015; Wolff et al., 2017). Because the star was presented on a random jitter, we realigned the data to the star onset at 0 seconds. Whereas PC1 in the star-delay reflected a dominant WM effect throughout the epoch that was unaffected by the task-irrelevant stimulus (see Figure 3C), the other three PCs revealed latent WM representations.

#### 3.3.1 Star stimulus modulates DM processing and “pings” latent WM representations

PC2 demonstrated that introducing the star fundamentally altered the temporal relationship between WM- and DM-related activity during the delay. Before onset of the star, PC2 captured an early test greater than sample effect which was significant for about 475 ms (-0.750- -0.275 s from star onset, *p* = 0.002; Figure 6A). Following this DM effect, test activity decreased and inverted with sample activity about 200ms before star onset, with sample activity gradually climbing in transition to a significant WM effect that began shortly after offset of the star and was sustained through the remainder of the delay (0.135-0.850 s from star onset, *p* = 0.002). PC2 explained 14.02% of variance in the star-delay period, and as with other PCs containing significant DM effects, PC2-loading electrodes were highly sampled from IFG (10 electrodes, 14.1%), SMG (7 electrodes, 9.0%), and AG (7 electrodes, 11.5%). Additionally, the ratio of dorsal frontal PC2-loading electrodes was more in SFG (SFG: 13 electrodes; MFG 19 electrodes; 0.73) than in PC1-loading electrodes (SFG: 17 electrodes; MFG: 31 electrodes, 0.55). Relative to the observed DM effect in PC2 of the uninterrupted delay period (see Figure 5A), the significant DM effect emerged earlier during the delay period and exhibited a declining rather than ramping trajectory. This shift suggests that introducing the star altered the natural evolution of the deliberative DM process. Of note, the star was always presented at the same jittered time during both sample and test delay periods within each star trial to control for visual processing demands. As a result, subjects were able to anticipate presentation of the star during test but not sample. This may explain why decreases in test activity began prior to star onset, in expectation of the interruption, and may also result in the deliberative DM process occurring earlier in the delay period in order to compensate for the interruption.

**Figure 6.**
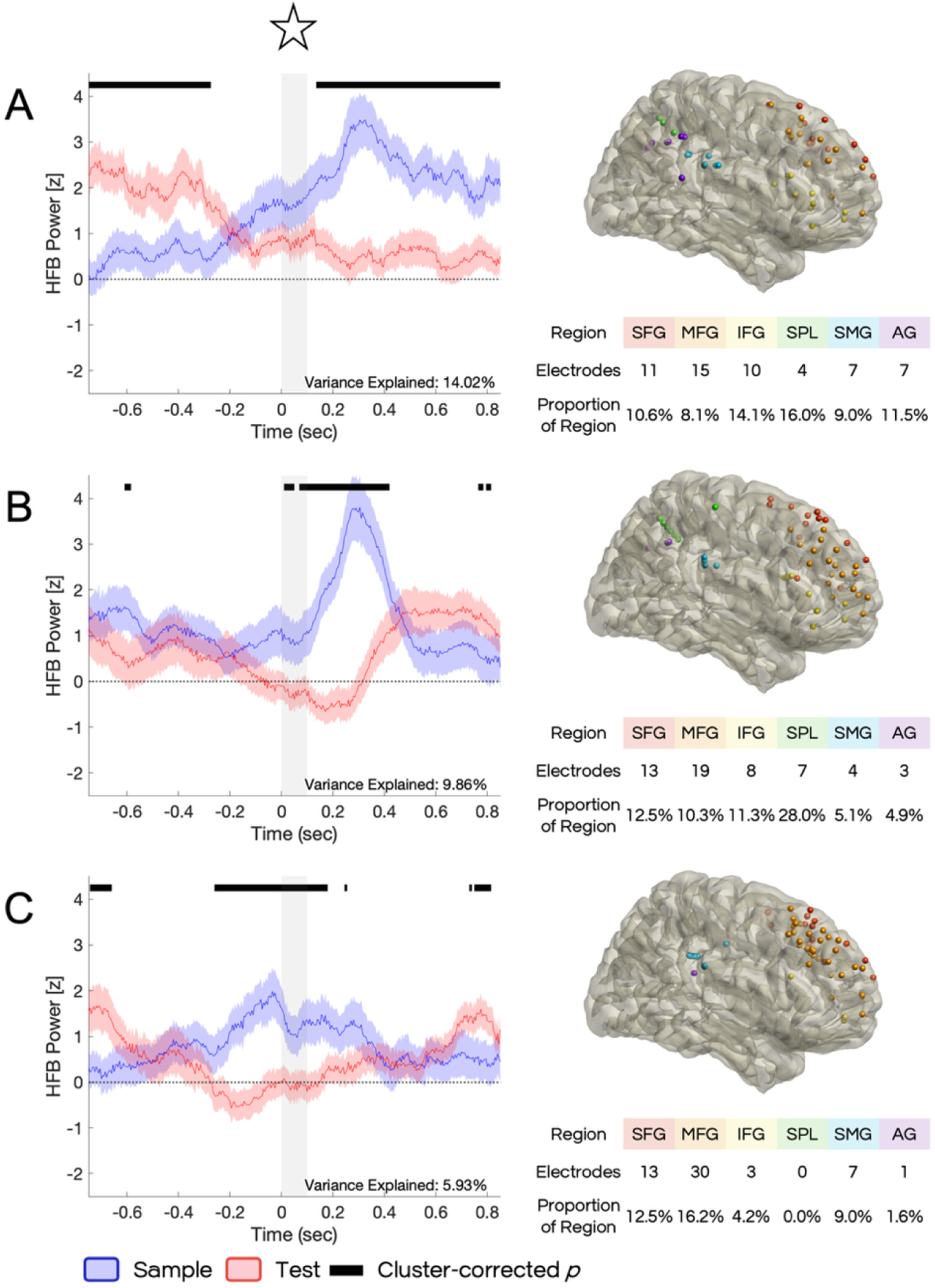
Working memory effects associated with interrupting the delay period, star delay PC2-4. Mean HFB timeseries from and locations of electrodes highly loading (75^th^ percentile of loadings) into the second (A), third (B), and fourth (C) PCs of the star delay. Left: Mean z-scored HFB traces, with standard error shown in shaded area, for sample (blue) vs. test (red). Black horizontal line indicates time points of significance by cluster-corrected *p*-value. Vertical grey shading represents duration of stimuli, with relevant stimulus represented above. Right, above: PC-loading electrodes shown on an MNI template brain, in lateral perspective and mirrored across hemispheres. Right, below: breakdown of electrodes by subregion, by number and relative sampling compared to cohort electrode coverage in each region.

The appearance of a significant WM effect following the star during sample delay, where it did not appear before, confirms that the star successfully performed its intended effect of eliciting measurable responses related to otherwise latent WM representations (Wolff et al., 2015). It is possible that the reactivation of WM-related processing during the sample delay facilitated subsequent evaluation of the test sequence and enabled the earlier emergence of DM-related activity during the test delay before the star (despite the effect attenuating once the star was presented).

#### 3.3.2 Novelty of star stimulus

Whereas PC2 reflected perturbation of ongoing DM processes by introducing the star, PC3 and PC4 primarily captured transient responses synchronized with the presentation of the star stimulus itself. PC3 isolated the strongest star-evoked response. Sample activity increased for about 410 ms after the star before returning to pre-star levels (Figure 6B) and was elevated compared to test beginning at the onset of the star (0.010-0.420 s, *p* = 0.002). This PC explained 9.86% of variance in the star-delay. As with stimulus-associated effects during the stimulus presentation period (see Figure 4), PC3-loading electrodes contained a larger proportion of electrodes from the SPL (7 electrodes, 28.0%) than non-stimulus-linked PCs. Star-associated effects were also observed in PC4, which explained 5.93% of variance. PC4 similarly captured a transient WM response associated with the star, although this response peaked immediately before rather than after star onset (Figure 6C), and was elevated compared to test activity for about 515 ms (-0.260-0.255 s from star onset, *p* = 0.018). Additionally, no PC4-loading electrodes were located in SPL, instead being in greater proportion from MFG (30 electrodes, 16.2%) and SMG (7 electrodes, 9.0%) than PC3 (MFG: 19 electrodes, 10.3%; SMG: 4 electrodes, 5.1%).

Both PCs also contained brief transient DM effects, occurring both before and after the star-associated WM effect. In PC3, these lasted for 25 ms (-0.610- -0.585 s, *p* = 0.002) and 50 ms (0.765-0.815 s, *p =* 0.002), and in PC4 both lasted 85 ms (-0.745 - - 0.660 s, *p* = 0.002; 0.730-0.815 s, *p* = 0.018). Together, these findings indicate that the star elicited two separable neural responses: reactivation of latent WM representations (captured primarily by PC2) and transient stimulus-evoked activity associated with the unexpected visual event (captured by PC3 and PC4). Both responses were accompanied by attenuation of ongoing DM-related activity during the test delay.

### 3.4 Varied anatomical distribution of electrodes

The PCA results revealed recurring trends in the anatomical distribution of the PC-loading electrode sets, including preferential recruitment of IFG in DM-related components and differences among parietal subregions, suggesting that distributed WM and DM processes contain anatomically specialized nodes within frontoparietal cortices. The identified WM and DM functions were represented across frontoparietal cortices, but with differing contributions from each area depending on the function, and all PCs across all trial segments included both frontal and parietal electrodes. This is suggestive of functional networks or distributed local processing underlying these cognitive processes. However, the trends in the distribution of the electrodes may point to critical nodes or anatomical organization within these networks. To test the hypothesis that subregions of the frontal and parietal cortices are functional specialized, we compared sample and test HFB activity within anatomically defined subregions in each trial segment.

#### 3.4.1 Functional gradient in prefrontal cortex

The PCA suggested a dorsal-ventral functional gradient within PFC, with WM-related components preferentially sampling dorsal frontal regions and DM-related components recruiting IFG. To directly test the hypothesis that frontal DM functions would be focally ventral to WM function, we compared sample and test HFB activity within anatomically defined IFG, MFG, and SFG.

IFG exhibited robust DM-related activity throughout the task but no WM effects. Significant DM effects were prevalent during both the stimulus presentation (0.020-0.175 s from shape 1 onset, *p* = 0.008; 0.450-1.005s, *p* = 0.006) and no-star delay (1.945-2.700 s, *p* = 0.018) periods (Figure 7A). Furthermore, as in PC2 of the star delay, onset of the star attenuated a significant and sustained DM effect (-0.740-0.000 s from star onset, *p* = 0.018), returning test activity to baseline (Figure 7A, right). The increase in sample activity observed in star-delay PC2 was not apparent, nor were any WM effects observed in IFG at any point in the trial. Together, these findings support the data-driven hypothesis that IFG is preferentially associated with DM-related processing, and highlight IFG as the only lateral frontoparietal subregion uniquely associated with either WM or DM.

**Figure 7.**
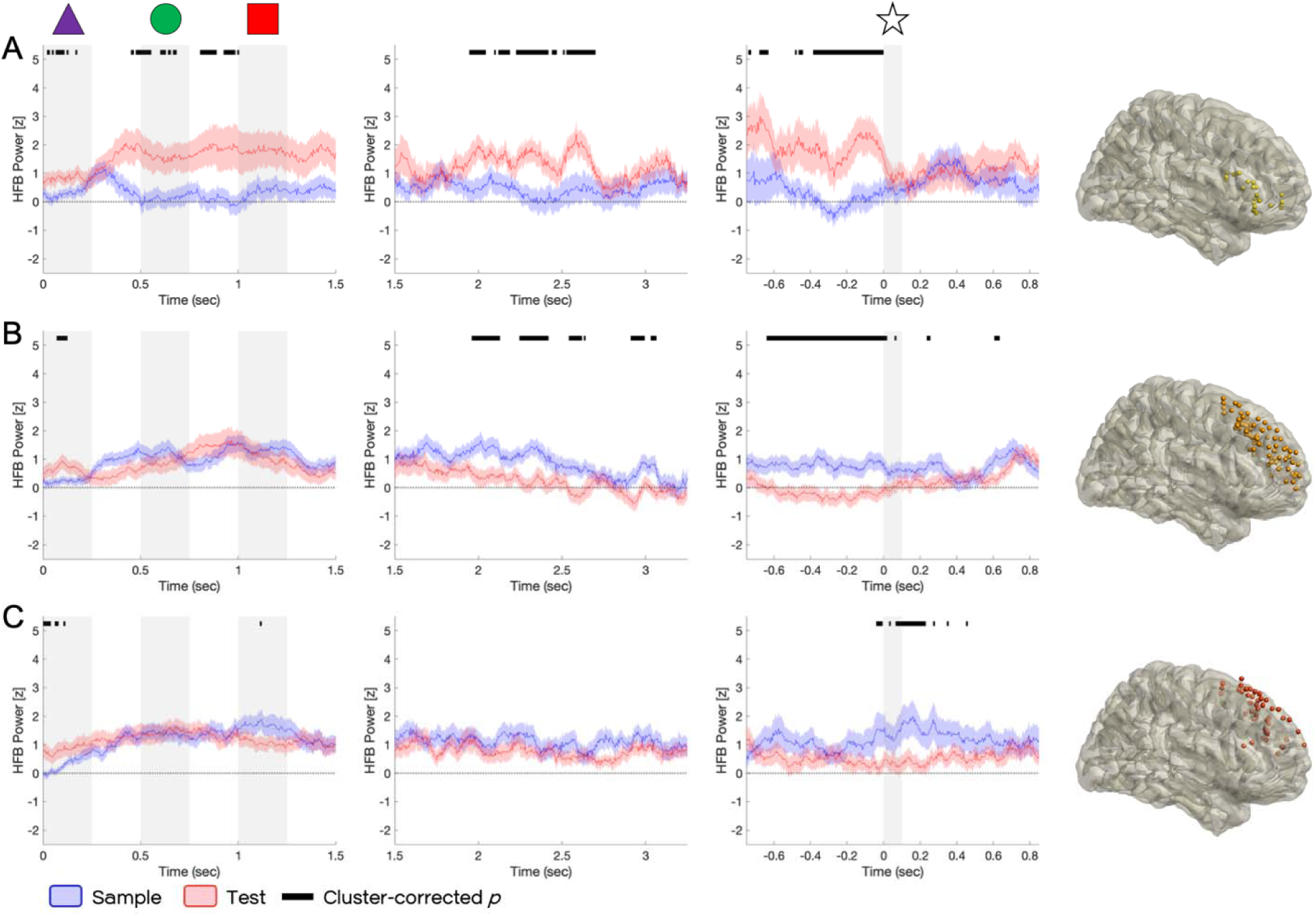
Effects localized to frontal subregions across trial segments. Mean HFB timeseries from electrodes in the inferior (A), middle (B), and superior lateral prefrontal gyri (C) during each trial segment: stimulus presentation, left; no-star delay, middle; and star delay, right. Mean z-scored HFB traces, with standard error shown in shaded area, for sample (blue) vs. test (red). Black horizontal line indicates time points of significance by cluster-corrected *p*-value. Vertical grey shading represents duration of stimuli, with relevant stimulus represented above.

In contrast, our PCA results suggest that SFG and MFG should be associated with WM-related activity, particularly in sustained WM representations (see Figure 3). We observed a greater ratio of SFG to MFG electrodes when latent WM representations were reactivated in star-delay PC2 (Figure 6A) versus the sustained activity in PC1 (Figure 3C) and hypothesized that SFG would play a more dominant role in “activity silent” WM maintenance. In this anatomical analysis, SFG and MFG exhibited WM-related activity with elevated sample activity above baseline in SFG and MFG, but not IFG, electrodes (Figure 7). Furthermore, SFG and MFG electrodes reacted differently to the presentation of the star. MFG showed a sustained WM effect through the delay that disappeared with the presentation of the star (-0.640-0.020 s from star onset, *p* = 0.002; Figure 7B, right), while a significant WM effect in SFG emerged with presentation of the star (-0.040-0.460 s, *p* = 0.018; Figure 7C, right) as if “pinged” by the stimulus (Wolff et al., 2015). These results suggest that MFG areas drive active WM representations, while SFG areas are more responsible for storing latent representations.

Together, these results support the dorsal-ventral functional gradient within PFC suggested by our PCA findings, with ventrolateral PFC (IFG) preferentially associated with DM-related activity and dorsolateral frontal regions (MFG and SFG) preferentially associated with WM-related activity. Furthermore, the differential responses of SFG and MFG to the star suggest that distinct forms of WM representation are distributed across dorsolateral PFC, with latent representations more evident in SFG and active, persistent representations more evident in MFG.

#### 3.4.2 Stimulus-associated effects in posterior parietal cortex

The PCA result also suggested functional differentiation within PPC, with inferior parietal regions (SMG and AG) contributing preferentially to stimulus-related WM effects and SPL contributing to a distinct set of transient responses. To test these observations directly, we compared sample and test HFB activity within anatomically defined SMG, AG, and SPL electrode groups (Figure 8).

**Figure 8.**
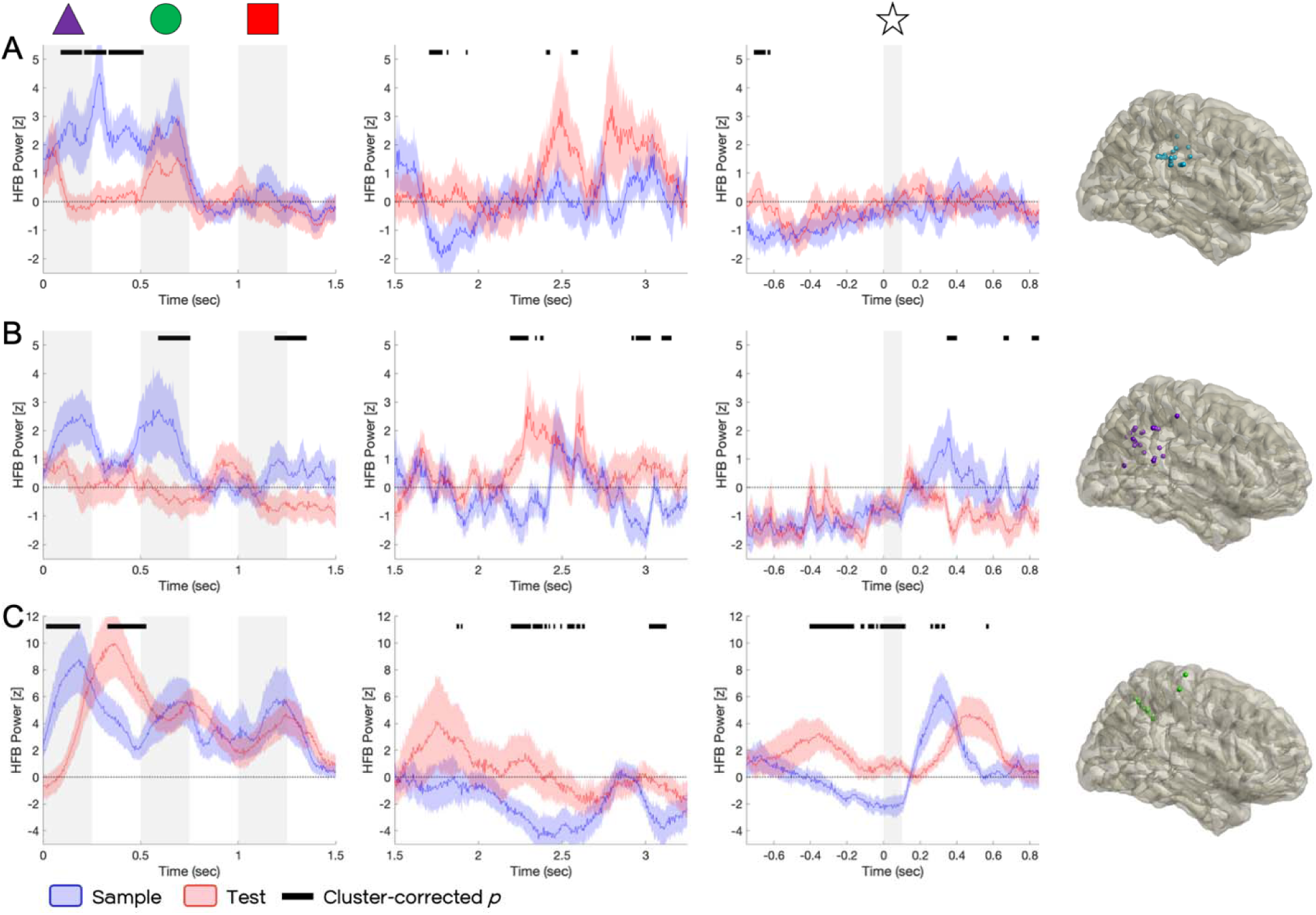
Effects localized to parietal subregions across trial segments. Mean HFB timeseries from electrodes in the supramarginal gyrus (A), angular gyrus (B), and superior parietal lobule (C) during each trial segment: stimulus presentation, left; no-star delay, middle; and star delay, right. Mean z-scored HFB traces, with standard error shown in shaded area, for sample (blue) vs. test (red). Black horizontal line indicates time points of significance by cluster-corrected *p*-value. Vertical grey shading represents duration of stimuli, with relevant stimulus represented above.

In anatomical analysis, SMG and AG subregions drove stimulus-specific WM effects, with activity patterns that closely resemble many of those extracted by PCA. In SMG, sample activity was significant during presentation of the first shape and the subsequent ISI (0.090-0.515 s from shape 1 onset, *p* = 0.002; Figure 8A, left), closely resembling the early stimulus-related WM effect captured by PC2 of stimulus presentation (see Figure 4A). Sample activity remained elevated through shape 2 before rapidly returning to baseline (also as in PC2). In AG, sample activity was significant during presentation of shape 2 (0.590-0.755 s from shape 1 onset, *p* = 0.002; Figure 8B, left), mirroring the shape 2-specific WM effect captured by PC4 (see Figure 4C). Together, these findings in SMG and AG indicate that the WM effects identified by the PCA were concentrated within inferior parietal cortex and that these regions are associated with detecting behaviorally relevant stimuli.

In contrast to SMG and AG, SPL exhibited a distinct temporal profile that closely resembled PC3 during stimulus presentation, the component which most highly sampled SPL electrodes (see Figure 4B). The SPL showed a significant WM effect during shape 1 (0.015-0.190 s from shape 1 onset, *p* = 0.002), followed by a significant test effect during the first ISI (0.330-0.530 s, *p* = 0.002; Figure 8C, left), reproducing the alternating sample- and test-related effects observed in PC3. Additionally, SPL exhibited an acute increase in sample activity following presentation of the star (0.255-0.335 s from star onset, *p* = 0.004, Figure 8C, right), similar to the transient star-evoked response observed in star-delay PC3 (see Figure 6B). These findings suggest that IPL inferior and superior parietal cortex contributed to distinct aspects of task processing: inferior parietal regions preferentially tracked behaviorally relevant stimuli for WM- and DM-related processing, whereas SPL generated brief responses associated with stimulus onset itself.

## 4. Discussion

WM and DM emerged as interacting processes that evolved across the trial to transform maintained representations into goal-directed decisions in contrast to distinct cognitive processes (Figure 9). HFB activity during a DMS task identified multiple temporally distinct patterns of neural activity associated with a dominant persistent WM representation, early stimulus-linked WM and DM processes organized around behavioral relevance, temporally separable early and late stages of decision formation, and latent WM representations that could be reactivated by perturbation. Although these processes were broadly distributed across frontal and parietal regions, they exhibited systematic differences in anatomical organization, including a dorsal-ventral functional gradient within PFC and differentiation among parietal subregions. Together, the results suggest that frontoparietal activity reflects a dynamic progression from memory encoding and maintenance to stimulus evaluation and decision formation, with WM and DM processes interacting throughout the trial as task demands evolve.

**Figure 9.**
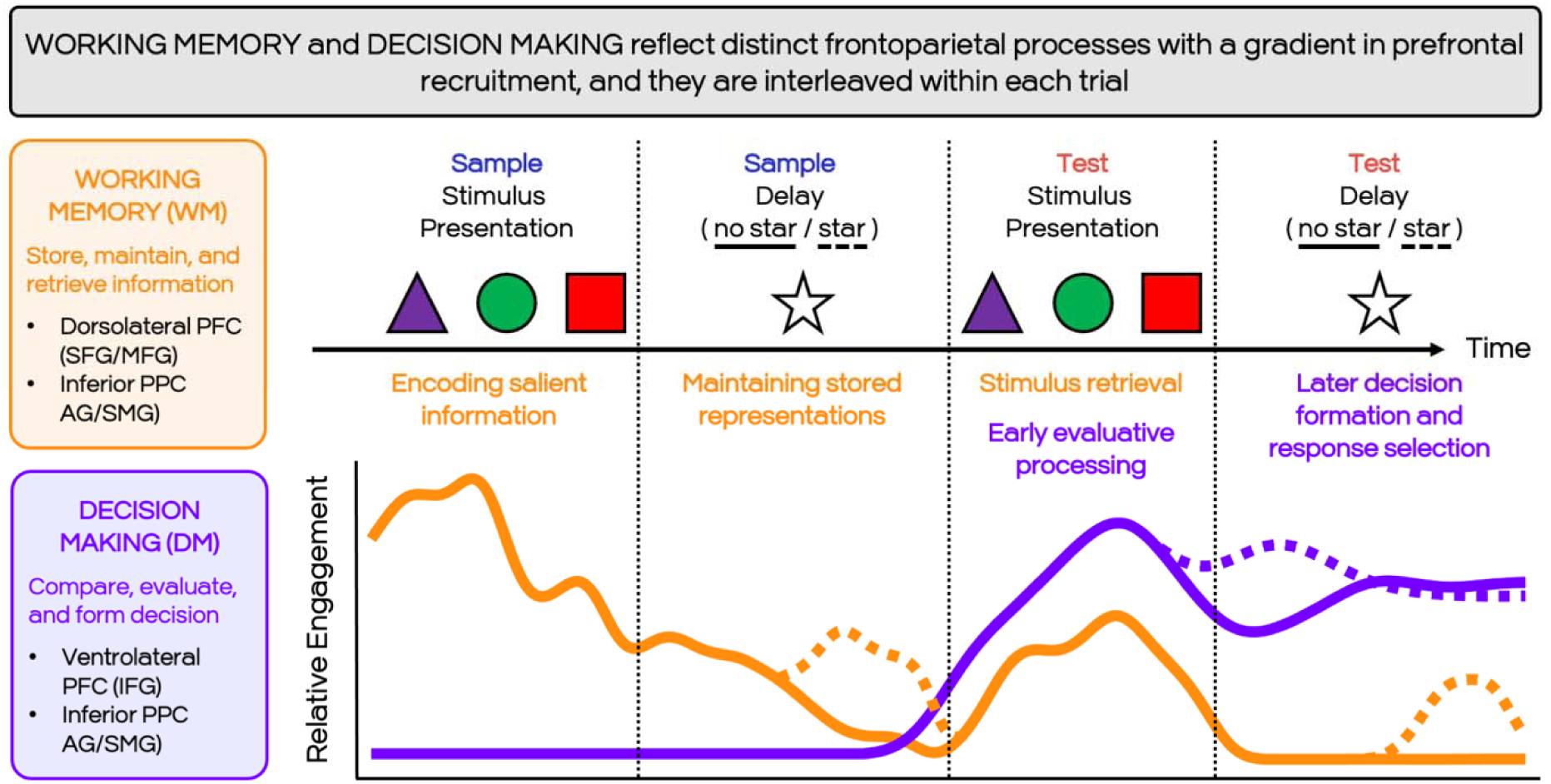
Conceptual schematic illustrating the temporal relationship between WM and DM processes during DMS performance. Solid lines summarize the evolution of WM (orange) and DM (purple) processes, during trials without a delay-period perturbation (solid lines) versus the effects of introducing the star stimulus during the delay (dashed lines). WM and DM are shown as separable but temporally interleaved processes that coexist within distributed frontoparietal networks rather than representing mutually exclusive brain states. Curves are schematic and intended to summarize the principal findings rather than depict the measured neural signals.

### 4.1 Persistent WM representation across trial phases

The most dominant pattern of neural activity throughout the task was an active, sustained WM-related process. Across all trial phases, the trajectory of the three PC1s converged on elevated sample-related HFB activity that persisted through stimulus presentation, delay, and star-interrupted delay epochs and was largely unaffected by perturbation. This stable component may reflect a persistent internal representation of the sample maintained across frontoparietal regions during task performance, and is consistent with classical models of WM in which information is maintained through sustained neural firing within frontoparietal circuits (Goldman-Rakic, 1995; Kamiński & Rutishauser, 2020; Miller & Cohen, 2001). This process was broadly distributed across frontal and parietal cortices, supporting the view that WM maintenance is implemented through distributed cortical networks. However, the strong representation in dorsolateral frontal regions suggests that MFG and SFG play a distinct role in sustaining actively maintained representations over time.

### 4.2 Early stimulus-linked processes reveal goal-driven, separable WM and DM dynamics

Of the observed stimulus-linked processes, the most dominant was organized around behavioral relevance rather than individual stimuli. While lower-variance components in stimulus presentation revealed activity associated with individual stimuli, the higher-variance PC2 distinguished between the behaviorally informative portion of the sequence (shapes 1 and 2) and the superfluous third shape. Because sufficient information to identify a mismatch was always available by the second shape, the restriction of these effects to the “salient” portion of the sequence suggests that WM encoding processes preferentially represented the information most informative for upcoming decisions rather than holding all presented stimuli equally.

DM-related activity exhibited a complementary pattern. In PC2, test activity increased with each successive stimulus, consistent with an accumulative decision process integrating information over time peaking around the onset of the third stimulus and then declining. This pattern suggests that evaluative processes began while the sequence was still unfolding and were primarily driven by behaviorally informative stimuli, dynamically allocating neural resources toward the information that contributes most directly to task performance.

The anatomical distribution of these effects further supports this interpretation. PC2-loading electrodes were enhanced in IFG and the inferior parietal lobe (SMG and AG), suggesting these regions enable salience detection in this task context. Previous studies have implicated them in prioritization of behaviorally relevant information and adaptive control (Erez & Duncan, 2015; Kahnt et al., 2014; Numssen et al., 2021; Singh-Curry & Husain, 2009). Our results suggest that these areas are recruited during both WM- and DM-driven processes to guide goal-oriented stimulus processing.

### 4.3 Decision-making processes continue during the delay period

Decision-related processing did not end with early evaluation during stimulus presentation. Instead, we observed a second stage of prolonged decision-related dynamics during the delay period. In PC2 of the uninterrupted delay, declining WM-related activity during the sample phase was accompanied by progressively increasing DM-related activity during the test phase. Viewed across the full trial, these trajectories suggest a natural progression from memory maintenance before presentation of the test sequence toward increasingly deliberative decision-related processing as subjects approached the behavioral response.

This later DM process differed qualitatively from the stimulus-linked effects observed during sequence presentation. Whereas early DM activity tracked important incoming evidence, delay-period activity exhibited a gradual ramping profile potentially reflecting processes involved in consolidating the decision or preparing for the upcoming response. Similar multi-stage decision processes have been proposed in evidence accumulation models in which early stimulus evaluation is followed by additional evidence accumulation prior to response execution (Brosnan et al., 2020; Liu & Pleskac, 2011; Pedersen et al., 2015; Usher et al., 2013). The presence of both early and late decision-related signals indicates that DM processes unfold across multiple timescales during WM tasks.

### 4.4 Star perturbation reveals latent WM representations and modified DM processes

The introduction of the task-irrelevant star stimulus provided a unique opportunity to probe the stability of ongoing neural processes during the delay. Consistent with previous work using perturbation approaches (Wolff et al., 2015), the star elicited transient increases in WM-related activity in all three of PC2-4, revealing neural responses associated with otherwise latent representations. This “pinging” effect suggests that at least some WM representations during the delay were maintained in a latent state through neural changes rather than active firing, in addition to the persistent representations captured in PC1, and that these active and activity-silent representations appear to coexist.

The emergence of the latent WM response was followed by an earlier DM effect than observed in the uninterrupted condition. Because participants could anticipate the upcoming interruption during the test delay (having encountered the star during the preceding sample delay), decision-related processing may have been initiated earlier to compensate for an expected interruption. Consistent with this interpretation, presentation of the star attenuated the ongoing DM effect, suggesting that the interruption temporarily disrupted deliberative processing. This suggests that the DM processes captured across frontoparietal regions are dynamic and flexible to adapt to changing cognitive demands. These findings also indicate that WM and DM are dynamically interdependent processes: viewed across the full trial, reactivation of WM-related processing during the sample delay may have facilitated subsequent evaluation of the test sequence, contributing to the earlier emergence of DM-related activity later in the trial.

### 4.5 Functional gradient across prefrontal cortex

The anatomical distribution of PC-loading electrodes revealed a systematic dorsal-ventral functional gradient within the PFC. WM-related effects were most associated with electrodes located in dorsolateral PFC regions corresponding to the SFG and MFG, whereas DM-related activity was consistently associated with electrodes in the IFG. This dissociation suggests that dorsolateral PFC areas preferentially support maintenance of task-relevant representations, while ventrolateral PFC plays a specialized role in evaluative decision processes. This finding is broadly consistent with proposals that dorsolateral PFC regions support maintenance and manipulation of information, whereas ventrolateral PFC regions support stimulus evaluation and selection processes (Badre & Nee, 2018; Badre & Wagner, 2007; Goldman-Rakic, 1995; Miller & Cohen, 2001).

The distinction extended beyond WM versus DM. Within dorsolateral PFC, MFG and SFG exhibited different responses to perturbation. MFG was associated with sustained WM activity that diminished following presentation of the star, whereas SFG exhibited WM-related activity that emerged only after perturbation. This pattern suggests that distinct forms of WM representation are distributed across dorsolateral PFC, with MFG contributing more to active maintenance and SFG contributing more to latent representations that can be reactivated when needed. This speculative interpretation provides a potential anatomical substrate for the coexistence of persistent and activity-silent WM states, a possibility raised in discussion of WM models in recent years (Barbosa et al., 2020; De Pittà & Brunel, 2022; Kamiński & Rutishauser, 2020; Miller et al., 2018).

Together, the present findings extend this view by demonstrating that the dorsal-ventral gradient reflects preferential specialization within a distributed network, with IFG contributing most to evaluative processing while dorsolateral prefrontal regions support persistent and latent WM representations. Rather than operating independently, these regions interact dynamically across the trial as task demands shift from memory maintenance to decision formation.

### 4.6 Stimulus-linked processes in posterior parietal cortex

PPC regions also exhibited functional differentiation, showing activity patterns closely tied to stimulus presentation. Inferior parietal regions (SMG and AG) recapitulated the stimulus-linked WM effects identified by PCA and selectively tracked behaviorally informative stimuli. These findings suggest that inferior parietal cortex contributes preferentially to the encoding and prioritization of task-relevant information, consistent with previous reports that these areas play a role in detecting task-relevant salience (Erez & Duncan, 2015; Kahnt et al., 2014; Molenberghs et al., 2007; Numssen et al., 2021; Singh-Curry & Husain, 2009). In contrast, SPL activity tracked the onset of visual stimuli, both to each shape during stimulus presentation and the star stimulus during delay. Unlike SMG and AG, SPL responses were not selective for behaviorally relevant stimuli, instead occurring in both sample and test and not specific to behavioral relevance. This pattern could represent a transient attentional or visual processing signal associated with detecting external events, in line with previous characterization of SPL function (Ciaramelli et al., 2008; Molenberghs et al., 2007).

Together, these findings refine existing accounts of PPC function by demonstrating how distinct parietal subregions contribute to different aspects of WM-guided decision making within the same task. Our results suggest that these functions are expressed dynamically during task performance, with inferior parietal cortex preferentially tracking the behavioral significance of incoming stimuli and SPL tracking the appearance of new information that may require updating ongoing processing. This organization may allow frontoparietal networks to simultaneously evaluate whether incoming information is relevant for a decision while remaining sensitive to environmental events that necessitate a shift in cognitive state.

### 4.7 Limitations

Limitations of these findings need to be considered. Participants were patients with medically refractory epilepsy undergoing clinical monitoring. Although seizure-onset zones and artifact-contaminated electrodes were excluded and prior work suggests that interictal recordings provide valid measures of cognitive processing (Rossini et al., 2017), disease-related factors may still influence neural activity. Nonetheless, we performed rigorous artifact reduction and outlier rejection of iEEG data, and the observed behavioral performance was comparable to healthy controls, strengthening confidence that these factors did not drive the observed HFB patterns. Furthermore, electrode placement was solely determined by clinical need providing incomplete and nonuniform sampling of frontoparietal cortex, including relative under-sampling of parietal cortex. Although this sampling bias cannot be eliminated, our statistical approach was designed to mitigate its effect through the use of nested random effects in the linear mixed-effects models and orthogonal PCA, which helped account for variability in electrode coverage while identifying robust spatiotemporal activity patterns. Future research can address this gap by replicating our analyses in larger, more-densely sampled cohorts and extend our findings by expanding electrode selection beyond the anatomical constraints of frontoparietal cortices. Finally, interpretations regarding latent WM representations rely on indirect inference from perturbation-evoked responses. Although consistent with activity-silent accounts of WM (Wolff et al., 2015; Wolff et al., 2017), alternative mechanisms may also contribute to the observed effects.

### 4.8 Conclusions

Together, these findings demonstrate that WM and DM processes emerge from temporally structured interactions within distributed frontoparietal areas. By combining intracranial recordings with perturbation of the delay period, we reveal that WM representations exist simultaneously in both persistent and latent states and that these representations interact dynamically with decision-related computations across time. Early stimulus-related activity reflects selective encoding of behaviorally relevant stimuli and rapid decision computations, whereas later delay-period dynamics suggest continued deliberative evaluation prior to response. These results provide new insight into coordinated WM and DM processes during complex cognitive tasks. Future work integrating iEEG and perturbation approaches with neuroimaging and network-level analyses may further clarify how distributed cortical circuits coordinate memory maintenance, attentional selection, and decision formation during WM tasks.

## CReDiT Authorship Contribution Statement

**S.A.:** Conceptualization, Formal analysis, Funding acquisition, Investigation, Methodology, Visualization, Writing – original draft, Writing – review and editing. **S.G.:** Data curation, Project administration, Writing – review and editing. **A.D.:** Methodology, Software, Writing – review and editing. **D.K.S.:** Data curation, Resources. **K.D.:** Data curation, Resources. **I.S.:** Data curation, Resources. **F.G.:** Data curation, Resources. **S.S.:** Data curation, Methodology, Resources. **J.R.:** Data curation, Resources. **E.A.:** Data curation, Resources. **R.K.:** Conceptualization, Data curation, Funding acquisition, Project administration, Resources, Software, Writing – review and editing**. R.B.:** Investigation, Resources, Supervision, Writing – review and editing. **E.J.:** Conceptualization, Data Curation, Formal analysis, Funding acquisition, Investigation, Methodology, Project administration, Resources, Supervision, Software, Visualization, Writing – review and editing.

## Acknowledgements

The authors thank Jack J. Lin, Peter B. Weber, Edward F. Chang, and Kurtis I. Auguste for their contributions to data collection and initial preprocessing, as well as the members of the Dynamic Brain Lab and Human Cognitive Neuroscience Lab for their invaluable feedback. This research was supported in part through the computational resources and staff contributions provided for the Quest high performance computing facility at Northwestern University which is jointly supported by the Office of the Provost, the Office for Research, and Northwestern University Information Technology.

